# *Limosilactobacillus fermentum* ATG-V5 stimulates the immune response by enhancing gut microbiome and metabolite in a cyclophosphamide-induced immunosuppression mouse model

**DOI:** 10.1101/2024.04.25.591046

**Authors:** Arum Park, Yeon-Hwa Park, Yewon Kang, Juyi Park, Gun-Seok Park, Yo-Han Ju, Won Hwang, Jihee Kang

## Abstract

*Limosilactobacillus fermentum* is known to exert immunomodulatory effects. In this study, we elucidated the immunoenhancing effects and mechanisms of a novel strain of *Limosilactobacillus fermentum* ATG-V5. Our findings demonstrate that ATG-V5 regulates the activation and proliferation of natural killer (NK) and T cells. It induced activation effects such as increased nitric oxide (NO) production, phagocytic capacity, and cytokine generation in Raw264.7 cells, which were confirmed to occur through the ERK and pp65 pathways. Furthermore, using a cyclophosphamide (CPP) mouse model, we demonstrated the immunomodulatory efficacy of ATG-V5, including enhanced activation of NK cells, increased levels of antibodies and cytokine secretion, in mice with immunosuppression induced by CPP. Additionally, by analyzing the gut bacterial community in the mouse model, we confirmed the ability of ATG-V5 consumption to regulate the gut microbiota, thereby substantiating the potential of this strain as a promising immunoenhancing agent.

## 1. Introduction

The twenty-first century has witnessed a wave of severe infectious disease outbreaks, not least the COVID-19 pandemic, which has had a devastating impact on lives and livelihoods around the globe (Baker et al., 2022). Effective vaccines have been developed to prevent infection, but there is a risk of infection with viruses and pathogens that have not yet been identified. The answer to this problem may be that a healthy functional immune system is paramount (Iddir et al., 2020; Jahns et al., 2018). It can strengthen the immune system by enhancing the phagocytic ability of macrophages, enhancing the activity of natural killer (NK) cells, and inducing the production of mucosal secretory immunoglobulin (Ig) A (Gaforio, Ortega, Algarra, Serrano, & Alvarez de Cienfuegos, 2002; Hirayama, Iida, & Nakase, 2017; van Gool & van Egmond, 2020). Also, interactions between the commensal microbiome and mammalian immune system development and function include multifaceted interactions in homeostasis and disease (Zheng, Liwinski, & Elinav, 2020). To this end, many researchers are exploring substances with immune-enhancing properties and studying the mechanisms of these properties.

The intestinal microbiota has co-evolved with the immune system1 to maintain a symbiotic relationship and aids in the digestion of inaccessible nutrients, provides colonization resistance to pathogens, and educates the immune system (Belkaid & Hand, 2014). Dynamic crosstalk between the microbiota and the host immune system during compositional or metabolic changes in the microbiota is essential for maintaining local and systemic immune homeostasis (Jiao, Wu, Huntington, & Zhang, 2020; Wiertsema, van Bergenhenegouwen, Garssen, & Knippels, 2021). Dysbiosis of the gut microbiome is associated with various changes in the immune system. Several references have demonstrated a role for the symbiont in promoting the differentiation of RORγt+ NKp46+ NK1.1hi non-conventional NK cells that produce IL-22 but not IL-17. This may indicate intestinal dysbiosis could potentially interfere with IL-22 and IL-17 production (Jiao et al., 2020; Sanos et al., 2009). Microbial colonization profoundly affects the intestinal CD4+ T-cell compartment and provides signals that differentiate naive T cells into Tbet+ Th1, GATA3+ Th2, RORγt+ Th17, FOXP3+ Tregs and BCL-6+ T follicle helper cells (Tfh) (Geuking & Burkhard, 2020; Littman & Rudensky, 2010). The microbiota can modulate the immune system by activating the AhR transcription factor. AhR ligands can be derived from many sources. Besides xenobiotic compounds, such as dioxin, AhR also recognizes natural ligands derived from cruciferous vegetables, as well as ligands produced by the microbiota through tryptophan metabolism (Geuking & Burkhard, 2020; Shinde & McGaha, 2018). For example, *Lactobacillus reuteri* provides indole derivates, such as indole-3 lactic acid, of dietary tryptophan that activate AhR (Cervantes-Barragan et al., 2017).

Probiotics, as defined by the Food and Agriculture Organization (FAO) and the World Health Organization (WHO), are living microorganisms that, when ingested in sufficient quantities, provide health benefits to the host(Food, Nations, & Report, 2001). Their primary function is to enhance the intestinal environment by promoting beneficial bacteria growth while inhibiting harmful bacteria’s proliferation (F. Yan & Polk, 2011). The gut microbiome has been found to play an important role in regulating the immune response, and various health benefits such as strengthening immunity, controlling hypersensitivity (allergy), and suppressing inflammation are claimed (Darma et al., 2020; Mazziotta, Tognon, Martini, Torreggiani, & Rotondo, 2023; F. Yan & Polk, 2011; Zheng et al., 2020). It has been recognized that changes in gut microbiome-derived metabolites are associated with immune responses in many immune-related diseases (Hayase & Jenq, 2021; Strazar et al., 2021; Yang & Cong, 2021). These metabolites can be produced by the gut microbiome, either from dietary components or by the host, and can be modified by gut bacteria or synthesized de novo. Metabolites derived from the gut microbiome affect numerous immune cell responses including T cells, B cells, dendritic cells, and macrophages (Gaforio et al., 2002; Geuking & Burkhard, 2020).

Cyclophosphamide (CPP) is an alkylating agent belonging to the group of oxazaphosporines (Emadi, Jones, & Brodsky, 2009). As CPP has been in clinical use for more than 40 years, there is much experience using this drug for the treatment of cancer and as an immunosuppressive agent for the treatment of autoimmune and immune-mediated diseases (Bensinger et al., 2001; Passweg & Tyndall, 2007). Ying et al. 2020 showed that high doses of CPP could damage the gastric mucosa and induce changes in the gut microbiome composition, leading to the translocation of pathogenic bacteria to vital organs of the body (Ying et al., 2020). CPP has been shown to disrupt of the intestinal barrier and digestive system problems in cancer patients undergoing chemotherapy, suggesting an association between CPP and gut microbiota (van Vliet, Harmsen, de Bont, & Tissing, 2010; van Vliet et al., 2009). CPP is used to evaluate the protective and enhancement effects on immunosuppression of drugs that require efficacy evaluation by treating mice (H. Yan et al., 2021). In this paper, we confirmed the efficacy of ATG-V5 in CPP-induced immunocompromised mice to confirm the efficacy of ATG-V5, whose immune-enhancing effect was confirmed *in vitro*.

## 2. Materials and Methods

### 2.1 Preparation of L. fermentum ATG-V5

L. *fermentum* ATG-V5 isolated from Korean Citrus reticulata (Korean Collection for Type Culture [KCTC 14481BP]) was incubated in Lactobacilli MRS Broth (Difco, USA) at 37 ◦C for 24 h, then cell pellets were obtained by centrifugation (3000× g, 10 min, 4 ◦C), and washed three times with phosphate-buffered saline (PBS; pH 7.4, Welgene, Korea). To treat at *in vitro* assay and *in vivo* mouse model, the ATG-V5 was suspended in PBS, and this was serially diluted to 10^-9^. After a serial dilution, diluted ATG-V5 was inoculated onto MRS agar and incubated in 37 ◦C for 24 h. The colony number was counted, and the colony forming unit (CFU) was calculated by colony number x dilution factor.

### 2.2 Culture of Raw264.7 cells, Molt-4 cells, Yac-1 cells, and NK-92 cells

Macrophage cell line Raw264.7, T lymphoblast cell line Molt-4, and mouse lymphoma Yac-1 cell were purchased from KCLB (Korean Cell Line Back). The human NK cell line NK-92 was obtained from KRIBB (Korea Research Institute of Bioscience and Biotechnology). Raw264.7 cells were cultured in DMEM (Gibco, USA) supplemented with 10% FBS (Corning, USA), penicillin/streptomycin (final concentration, 100 U/ml, Gibco, USA). Molt-4 cells and yac-1 cells were cultured RPMI1640 (Gibco, USA) supplemented with 10% FBS, 1% penicillin/streptomycin. NK-92 cells were cultured MEM-a supplemented with 12.5% FBS, 12.5% Horse serum (Gibco, USA), 200 uM inositol (Sigma, USA), 100 uM 2-mercaptoethanol (Sigma, USA), 20 uM folic acid (Sigma, USA) and 20 ng/ml IL-2 (Peprotech, UA). All cells were cultured at 37 °C in a 5% CO_2_ incubator.

### 2.3 MTT assay in Raw264.7 cells and Molt-4 cells

To measure cell proliferation, Raw264.7 cells (1×10^5^ cells/well) or Molt-4 cells (1×10^5^ cells/well) were seeded into each 96-well plates in 100 μl of plain medium and incubated at 37 °C in a 5% CO_2_ incubator for 6 h. After 6 h of incubation, Raw264.7 cells or Molt-4 cells were treated with cell 1: ATG-V5 0, 5, 10, 20, 40 MOI rate (multiplicity of infection, cell:bacteria), and were incubated at 37 °C in 5% CO_2_ for 24 h. After 24 h of incubation, the absorbance was measured at 450 nm using tetrazolim salt WST-1 [2-(4-Iodophenyl)-3-(4-nitrophenyl)5-(2,4-disulfophenyl)-2H-tetrazolium]-based EZ-cytox solution (Daeillab, Korea) according to the manufacturer’s instructions. Briefly, 20 μl of EZ-cytox solution was added to each well followed by incubation at 37 °C in a 5% CO_2_ incubator for 1 h. The absorbance was measured immediately at 450 nm.

### 2.4 Phagocytosis of Raw264.7 cells

To measure cell proliferation, Raw264.7 cells (1×10^5^ cells/well) were seeded into a 96-well plate in 100 μl of plain medium and incubated at 37 °C in a 5% CO_2_ incubator for 24 h. After 24 h of incubation, Raw264.7 cells were treated with cell 1 : ATG-V5 0, 5, 10, 20, 40 MOI rate and were incubated at 37 °C in 5% CO_2_ for 1 h. The phagocytosis was measured at 405 nm using Phagocytosis Assay kit (with zymosan substrate) (Abcam, UK) according to the manufacturer’s instructions. Briefly, after washing with PBS at 3 times, Raw264.7 cells were incubated pre-labeled zymosan particles. After removing culture medium and washing, the Raw264.7 cells were added and incubated fixation solution for 5 min, blocking reagent for 60 min, permeabilization solution for 5 min, and detection reagent for 60 min. Washing was performed at each step. After washing, each well was incubated to detection buffer for 10 min and added substrate and incubated for 5-20 min. Subsequently, the Raw264.7 cells were treated with a stop solution, and the absorbance at 405 nm was promptly measured.

### 2.5 Nitric oxide (NO) secretion by Raw264.7 cells

Raw264.7 cells of 1×10^5^ cells/well were seeded into a 96-well culture plate. After 24 h, the cells were pre-incubated with different sample concentrations for 24 h, as mentioned above. LPS 1 µg/ml (Sigma, USA) was used as a positive control. The NO concentrations were estimated by calculating the quantity of released nitrite, using Griess reagent (Sigma, USA), in the cell supernatant. The OD was determined at 540 nm using a microplate reader.

### 2.6 Protein expression levels of p-p65, p-65, p-ERK, ERK, and GAPDH using western blotting

Raw264.7 cells were seeded in six-well plates at 5x10^5^ cells/well. After seeding, cells were starved overnight, treated with ATG-V5 (0, 5, 10, 20, 40 MOI rate) or LPS 1 ug/ml (positive control) for 1 h. Cells were treated with LIPA cell lysis buffer (Sigma, US) for 30 min on ice with occasional vortex. The cell lysates were resolved on 4 – 12% SDS-PAGE gels, followed by transfer to nitrocellulose membranes. The membranes were blocked in superblock solution at RT for 1h, and then incubated with the appropriate primary antibodies against p-p65 (Cell Signaling Technology, CST, US), p65, p-ERK, ERK, or GAPDH in superblock solution (Thermo Fisher Scientific, USA) at 4°C overnight. The membranes were washed (5 min, five times) and incubated for 1 h with HRP-conjugated secondary antibodies diluted to 1:5000. After washing, the membranes were incubated with ECL reagents (Thermo Fisher Scientific) and chemiluminescent signals were visualized using Chemidoc (Bio-rad, USA). Western blot bands from three experiments were quantitated by densitometric analysis.

### 2.7 Measurement of cytokines or Immunoglobulins

The levels of IL-6, TNF-α and IL-10 were determined using ELISA kits (Biolegend, USA), according to the manufacturer’s instructions. Briefly, the Raw264.7 cells were cultured for 24 h with different concentrations of the sample, as described above. LPS 1 µg/ml was used as a positive control. The supernatant was harvested and the number of cytokines was measured using the respective ELISA kit.

The serum was obtained by centrifuging the collected blood at 12000 rpm for 15 minutes. Mouse IFN-γ, IgG, IgA, and IgM ELISA kits were procured from Invitrogen (USA). ELISA plates (Nunc, Denmark) were coated with 1× mouse IgG, IgA, or IgM capture antibody in PBS at 4°C overnight. After coating, plates were washed with PBST (PBS containing 0.05% Tween-20) and blocked with PBS containing 1% BSA (Sigma) for 1 hour with shaking at room temperature (RT). After another washing with PBST, serum sample was diluted in PBSA (PBS containing 1% BSA) and added to each well. The plate was incubated at 4°C overnight with shaking. After washing, 1× HRP-conjugated detection antibody was added to each well and the plate was incubated at RT for 1 hour with shaking. After washing with PBST, TMB solution (SurModics, USA) was added to each well and the plate was incubated at RT with shaking. After incubation, stop solution (2N HCl) was added to each well. Optical density was then measured at 450 nm on a microplate reader (BioTek Instruments, Inc., USA).

### 2.8 NK cell or splenocytes-mediated cytotoxicity test

To measure NK-92 cells or splenocytes-mediated cytotoxicity, 50 μl of cultured NK-92 cells or splenocytes (5×10^5^ cells/well) and 50 μl of K562 or YAC-1 cells (1×10^4^ cells/well) were co-cultured in 96 -well for 4h in an CO_2_ incubator. The co-cultured cells were centrifuged at 2500 rpm for 5 minutes to precipitate the cells and 10 μl of the supernatant was transferred to a new 96-well plate. 100 μl of LDH reaction mixture (DoGenBio, Republic of Korea) was added to the supernatant and reacted at room temperature for 30 minutes under light blocking, and then the reaction was stopped by adding 10 μl of stop solution to each well. After the reaction was completed, the absorbance of each well was measured at 450 nm.

### 2.9 Animals

In this case, eight-week-old ICR male mice (Orientbio, Republic of Korea) weighing 35 ± 2 g were used. All mice were housed under controlled temperature (22 ± 2 ℃) and humidity (50–60%), with a 12 h light: 12 h dark cycle. The experimental protocol was approved by the Institutional Animal Care and Use Committee at the Atogen Co. Ltd. (No. ATG-IACUC-RDIM-201010).

### 2.10 Cyclophosphamide-induced immunosuppression mouse model

ICR mice were divided into four groups consisting of negative control (NC, PBS-treated group), CPP (Cyclophosphamide 150 mg/kg (CPP, Sigma, USA) + PBS-treated group), PC (CPP + Levamisole (50 mg/kg, Sigma, USA), ATG-V5 8 (CPP + 1 × 10^8^ CFU/mouse), ATG-V5 9 (CPP + 1 × 10^9^ CFU/mouse), ATG-V5 10 (CPP + 1 × 10^10^ CFU/mouse) (n = 9∼10). Mice of all groups, except the NC group, were intraperitoneally injected with CCP (150 mg/kg/day) twice, once on day 0 and again on day 2, for induction of immunosuppression. Mice in all groups were orally administered with the PBS, levamisole, or ATG-V5 once a day on days 0 to 10 (11 days). All mice were sacrificed on day 10. Spleen and thymus were harvested from all mouse groups and weighed. Spleens were used for H&E staining and analysis of splenocytes-mediated cytotoxicity and T cell subsets. Blood was used for cytokine and antibody analysis. In the cecum, the intestinal microbiome was analyzed using the NGS system, and the intestinal metabolite was measured using LC-MS.

### 2.11 Histology of spleen

Spleen were dissected and processed for paraffin embedding and serial sectioning. The histologic sections were deparaffinized, rehydrated and stained with H&E. The severity of white pulp atrophy in spleen was determined according to the following criteria: 0 = Normal, 1 = mild, 2 = moderate, 3 = severe. The white pulp atrophy score for each mouse was the mean of white pulp atrophy severity of five spleens.

### 2.12 Quantitative real-time polymerase chain reaction (qRT-PCR) assay

Total RNA was extracted from spleen using TRIzol® reagent (Thermo Fisher Scientific), and 2 μg of total RNA was reverse transcribed into first-strand cDNA by using a reverse transcription reagent kit (TOYOBO, Japan) according to the manufacturer’s protocol. qRT-PCR was performed using SYBR® Green real-time PCR kit (Thermo Fisher Scientific, USA) in a 20 µl reaction volume on the 7500 Fast RT-PCR (Applied Biosystems, USA). GAPDH was used as internal standard to normalize gene expression levels. The 2-ΔΔCT method was used to analyze relative gene expression. Each qRT-PCR experiment was replicated ≥ 3 times. Results are expressed as extent of change relative to control values. Primer that used to qPCR was showed on the table 1.

**Table 1.**
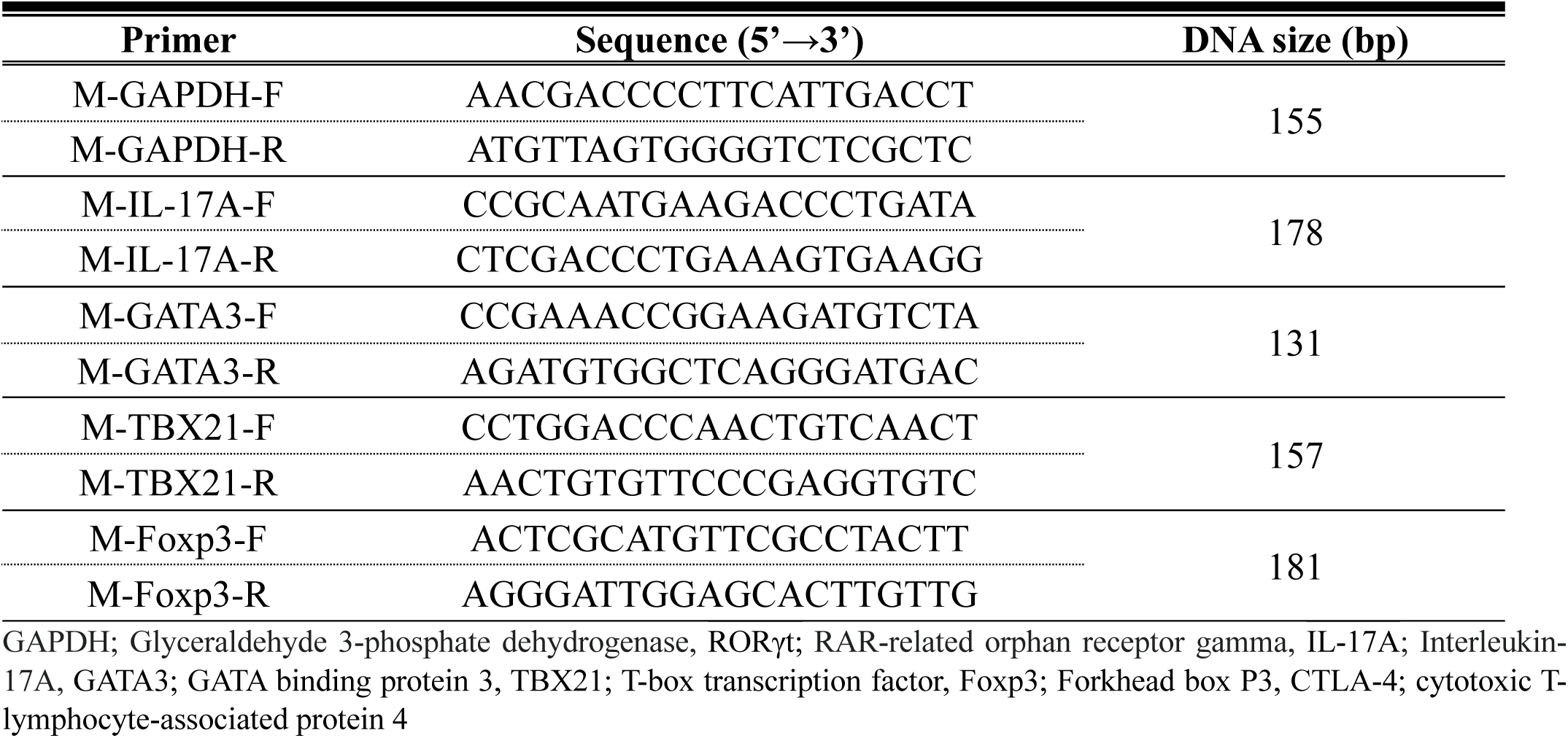
Primer list.

### 2.13 Caecal microbiota analysis

Genomic DNA was extracted from fecal samples using the QIAamp PowerFecal Pro DNA Kit (QIAGEN). The quantity and quality of extracted DNA were measured using a Qubit 3.0 Fluorometer (Thermo Fisher Scientific) and agarose gel electrophoresis, respectively. The V4 hypervariable regions of the bacterial 16S rRNA were amplified with unique 8 bp barcodes and sequenced on the Illumina MiSeq PE300 platform according to the standard protocol (Caporaso et al., 2012). Raw reads were analyzed using the QIIME2 pipeline (Bolyen et al., 2018) with the SILVA 138 database (Yilmaz et al., 2014). The non-parametric Kruskal-Wallis test was used to compare the differences in diversity indices and microbial taxa. The weighted UniFrac distances were previously obtained and used for PCoA (Lozupone, Lladser, Knights, Stombaugh, & Knight, 2010).

## 3. Results

### 3.1 Limosilactobacillus fermentum ATG-V5 enhances the activity of immune cells

To determine whether ATG-V5 has a potential effect on activity of immune cells, the macrophage cell line Raw264.7 was treated with ATG-V5 and then changes were confirmed. It was confirmed that the treatment of ATG-V5 did not affect the proliferation of Raw 264.7 cells (figure 1A). Treatment of Raw 264.7 cells with ATG-V5 increased phagocytosis (figure 1B), indicating that ATG-V5 stimulates of cellular immune responses. In addition, it was confirmed that nitric oxide was significantly produced in a concentration-dependent manner after treatment with ATG-V5 (figure 1C). The production of inflammatory cytokines IL-6 and TNF-a was significantly increased, and there was no significant difference in IL-10 (figure 1D). After treatment with ATG-V5, it was confirmed that ATG-V5 activates Raw264.7 cells through the activation of ERK and p65 by confirming the increase in the expression level of pp65 and pERK to confirm the activation pathway (figure E). Raw 264.7 cells are often used as macrophages to fight various bacteria and viruses. Therefore, the increased phagocytosis of Raw 264.7 cells through ATG-V5 treatment suggests that the activation of the immune system and the ability to eliminate intracellular pathogens were enhanced. Additionally, it was confirmed that ATG-V5 increased the proliferation of the T cell line and molt-4 cell line (figure 1F), in a dose-dependently, and increased the cytolytic activity of the NK92 cell line (figure 1G).

**Figure 1.**
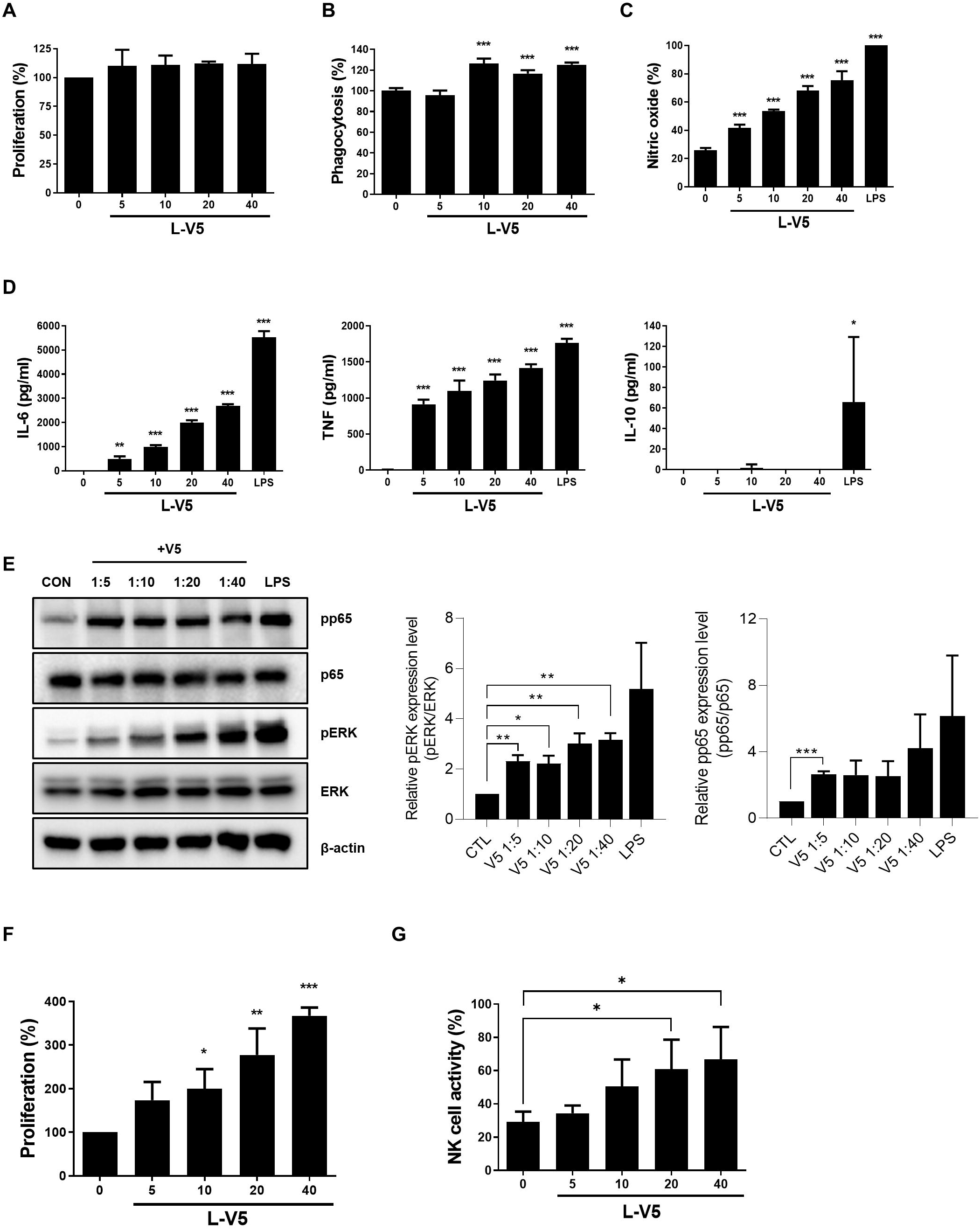
*Limosilactobacillus fermentum* ATG-V5 augments immune cell functionality. Raw264.7 cells were cultured for 24 hours in the absence or presence of various concentrations of *Limosilactobacillus fermentum* (ATG-V5) to determine (A) cell viability, (B) phagocytosis and (C) nitric oxide (NO) production. (D) Production amount of inflammatory cytokines according to the strain treatment was measured by ELISA. (E) Extracellular signal-regulated protein kinase (ERK) and nuclear factor-Κb(NF-Kb, p65) analyzed by Western blotting after the treatment of ATG-V5 in Raw 264.7 cells. β-actin served as a loading control. The graphs beside represent relative intensity defined as the intensity of the pp-65 or p-ERK normalized to ERK or p-65. (F) Proliferation results when Molt-4 was treated with ATG-V5 for 24 h by MTT assay. (G) NK92 cells were cultured in the absence or presence of ATG-V5 for 24h and cytotoxicity against K562 cells was examined by LDH assay. Bars represent mean ± SD. Statistical analyses were performed using the paired two-tailed Student t test. p value when compared with the control group (no treatment). *p < .05, **p < .01, and ***p < .001

### 3.2 Effects of Limosilactobacillus fermentum ATG-V5 on the size and weight of immune-related tissues

To evaluate the immune-activating efficacy of ATG-V5 in an *in vivo* system, we conducted experiments using a mouse model. CPP, an immunosuppressive agent, induced immune suppression in the mice. Subsequently, the mice were evaluated for immune response to ATG-V5. Injection of CPP into mice significantly increased spleen size and decreased thymus size (figure 2A). In contrast, the mice that received oral administration of ATG-V5 after injection of CPP showed a decrease in spleen size and weight, which had been increased by CPP, and recovery of thymus size and weight. These findings suggest that ATG-V5 may have a protective effect on immune organs in mice treated with CPP. In addition, we examined the white pulp of the spleen section. The spleen is divided into the red pulp, which filters red blood cells, and the white pulp, which activates immune responses through humoral and cell-mediated pathways. Atrophy of the white pulp occurs when the areas of T or B cells are lost. As a result, CPP treatment increased white pulp atrophy, and white pulp atrophy increased by CPP was significantly suppressed by ATG-V5 administration (figure 2B).

**Figure 2.**
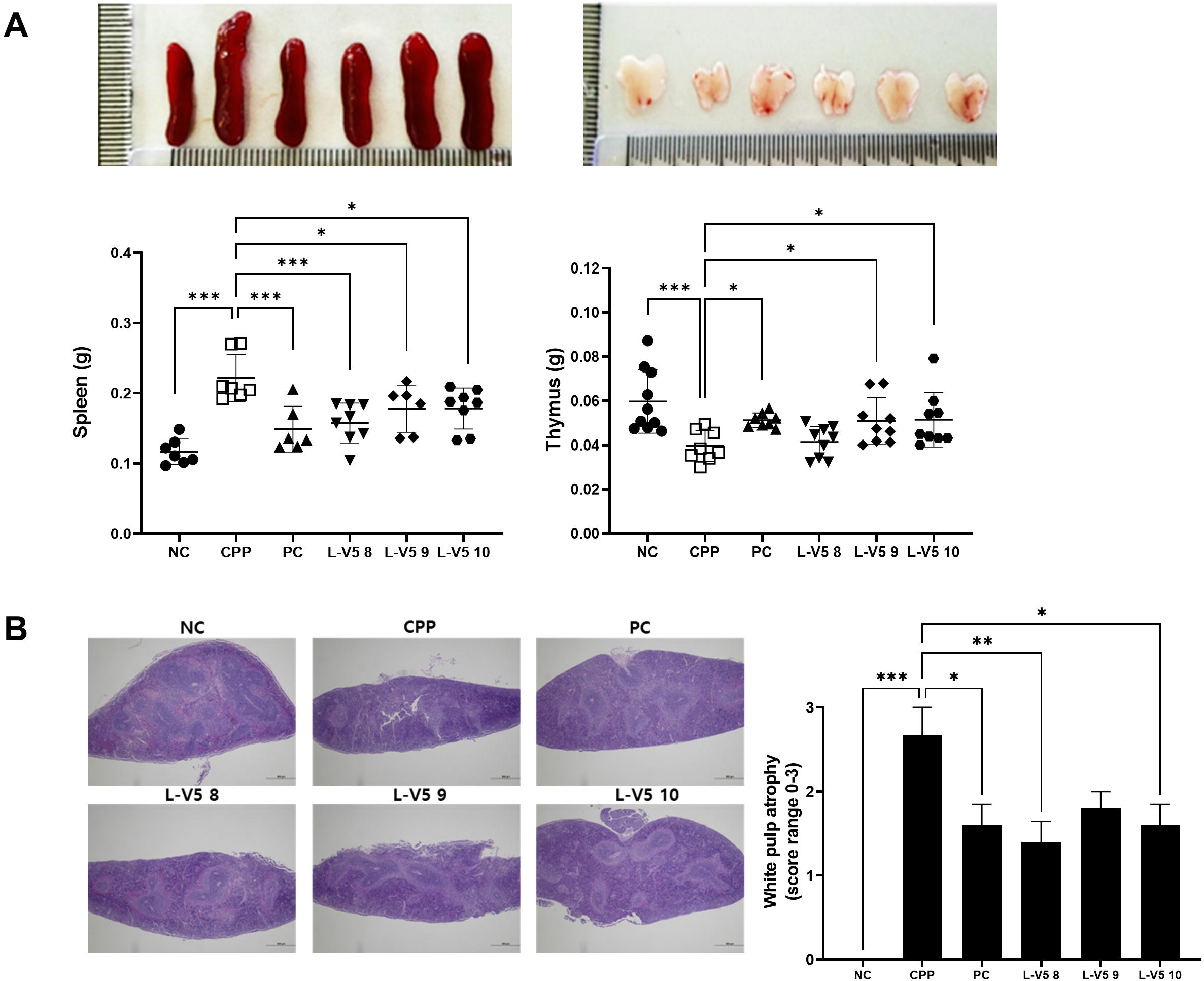
Effects of *Limosilactobacillus fermentum* ATG-V5 on immune-related tissue size and weight in CPP-induced model. (A) Comparison of the size and weight of the spleen and thymus in the control group (NC), the group treated with cyclophosphamide (CPP), and the group administrated the ATG-V5 after CPP treatment. (B) H&E stained sections of spleens were examined for microscopic changes. The severity of white pulp atrophy in spleen was determined according to the following criteria: 0 = Normal, 1 = mild, 2 = moderate, 3 = severe. The beside graph shows the average degree of white pulp atrophy for 5 spleens in each group. Bars represent mean ± SD. Statistical analyses were performed using the paired two-tailed Student t test. p value when compared with the CPP group. *p < .05, **p < .01, and ***p < .001

### 3.3 Limosilactobacillus fermentum ATG-V5 increases immune-enhancement in the CPP-induced immunodepression model

Several immune cells were analyzed as indicators to confirm that the immune capacity reduced by CPP in the mouse model was increased by ATG-V5. First, to confirm the change in the cytolytic activity of NK cells in the spleen, splenocytes were isolated and co-cultured with YAC-1, a target of NK cells. NK cell activity of splenocytes against Yac-1 cells in administering the ATG-V5 group was significantly increased compared to the CPP treatment group. Moreover, the increase between groups was concentration dependent on ATG-V5 (figure 3A). In addition, activated NK cells secrete IFN-γ to induce target cell death. As a result of the experiment, it was confirmed that blood IFN-γ decreased in the CPP treatment group and increased again in the AGT-V5 high-concentration administration group (figure 3B). Among immune cells, B cells induce a humoral immune response by producing antibodies called immunoglobulins (Ig) against specific antigens invaded from the outside. As a result of the experiment, it was confirmed that the concentrations of IgA, IgG, and IgM in the blood decreased by CPP treatment and increased in the ATG-V5 administration groups than in the CPP treatment group (figure 3C). In addition, the effect of ATG-V5 on CD4 T cell differentiation and immune response was confirmed. With mRNA isolated from splenocytes, changes in differentiation of T cell subsets, Th1, Th2, Th17, and Treg cells, were measured through the amount of mRNA expression. As a result of the experiment, the differentiation of naïve T cells was suppressed by CPP treatment, so the total number of T cell subsets was suppressed. Furthermore, it was confirmed that the number of Th1, Th2, Th17, and Treg cells increased significantly by taking ATG-V5(figure 3D).

**Figure 3.**
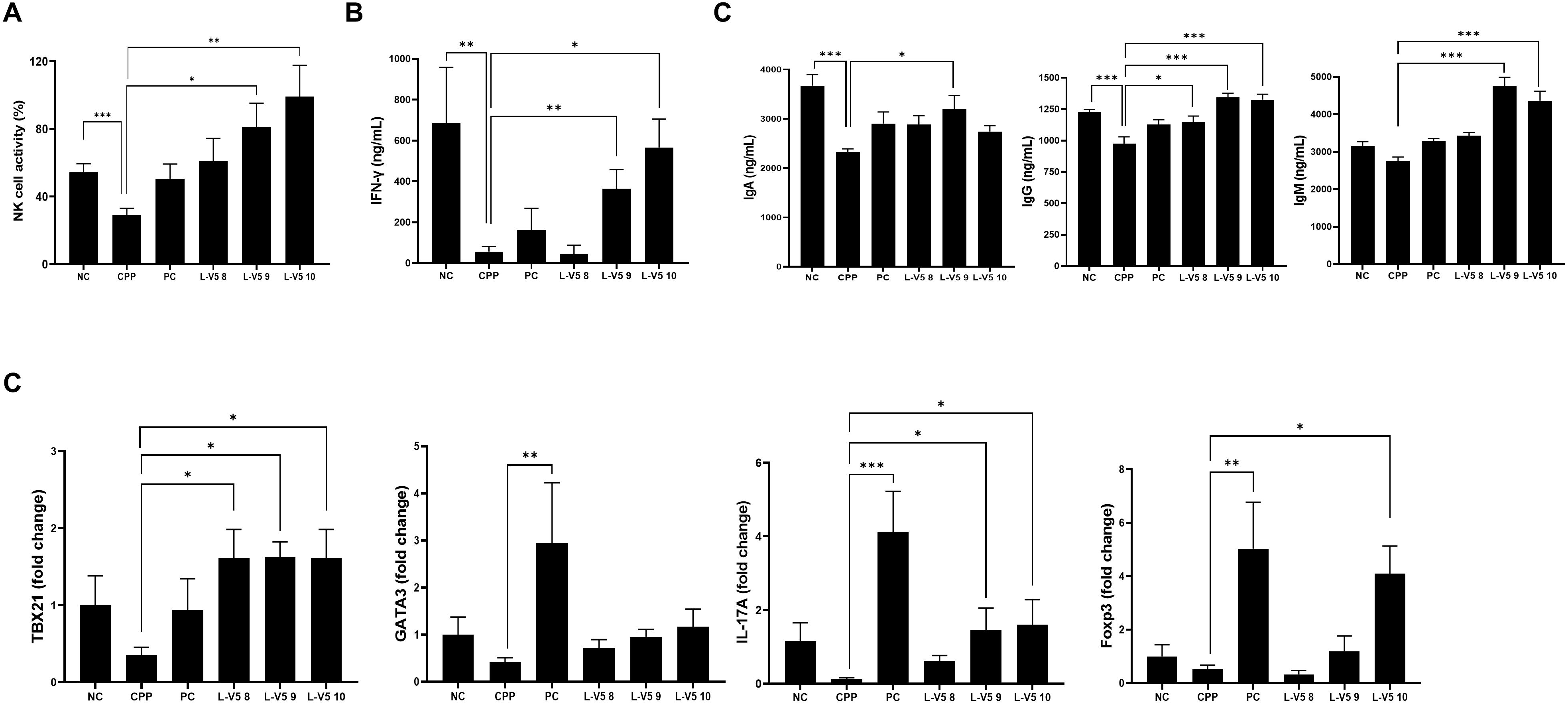
ATG-V5 induces immune enhancement in a CPP-induced model. (A) Mouse splenocytes from each group were isolated and NK cytotoxicity was measured against YAC-1 cells according to a 50:1 (splenocyte: YAC-1) ratio. The serum cytokine concentrations of (B) IFN-γ (C) IgA, IgG and IgM were detected by ELISA assay. (D) Expression of T cell subset related genes (TBX21, GATA3, IL-17A, Foxp3) was examined by RT-PCR. RNA levels are normalized to GAPDH gene. Bars represent mean ± SEM of three independent experiments. *p < .05, **p < .01, and ***p < .001

### 3.4 Modulation of gut microbiota by strain ATG-V5

To examine the effects of strain ATG-V5 on the gut microbiota, the bacterial community from the cecum samples of each experimental group was analyzed (figure 4A). Weighted UniFrac-based principal coordinate analysis (PCoA) was also performed to compare group differences in the overall microbiota profile (figure 4B). The 5 X 10^8^ ATG-V5-treated group and 5 X 10^9^ ATG-V5-treated group show relatively similar changes with the levamisole-treated group (positive control) compared with the 5 X 10^10^ ATG-V5-treated group. Bacterial richness was measured by Chao1, which means bacterial richness, and Shannon index, which means species diversity, showed that higher doses of ATG-V5-treated groups showed significantly different (figure 4C, D). At the genus level, the population of *Lactobacillus* was significantly decreased in the 5 X 10^10^ ATG-V5-treated group.

**Figure 4.**
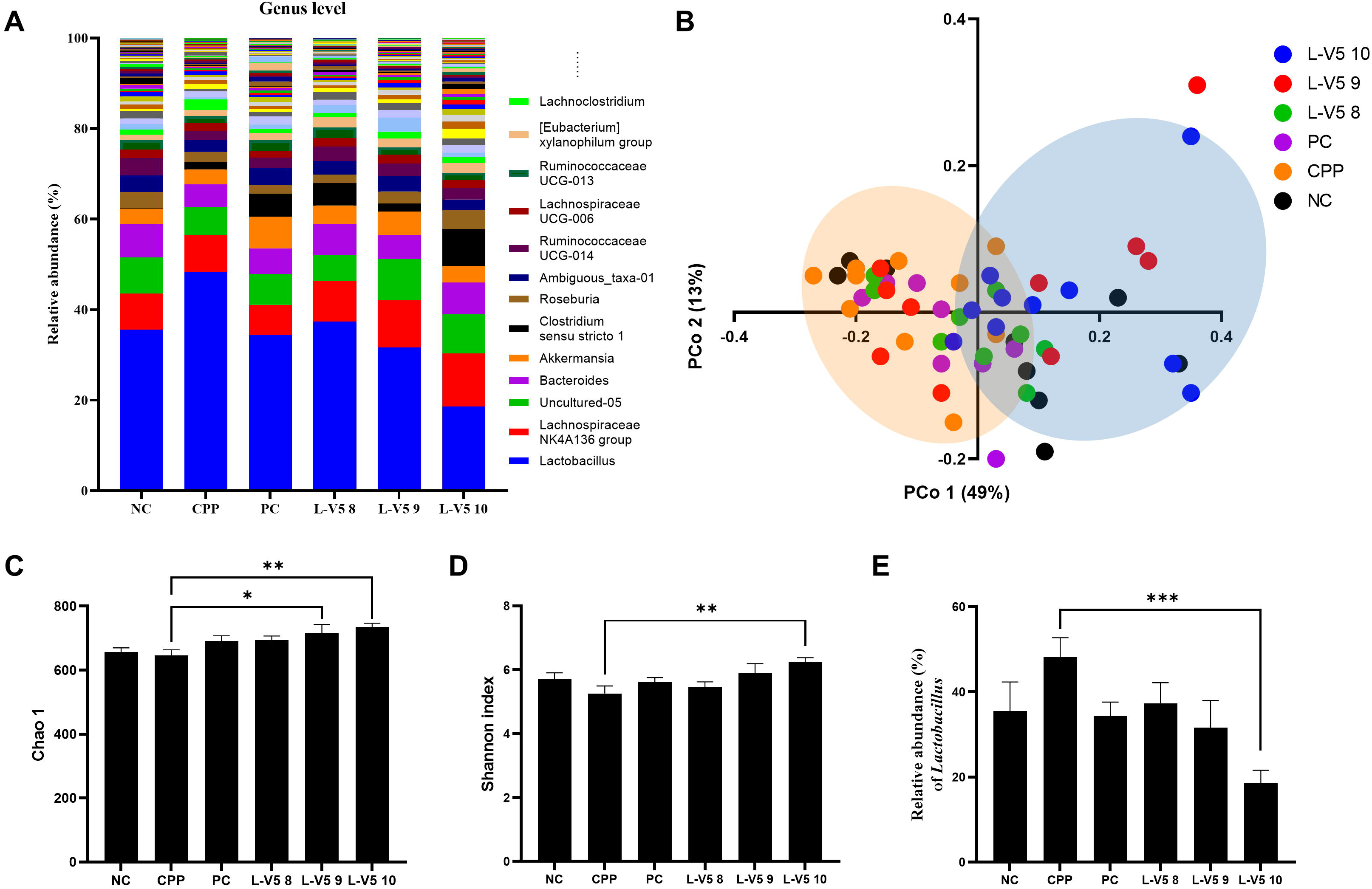
Change of caecal bacterial community by ATG-V5 treatment. (A) Average relative abundance of the genus level of taxonomy in each group. (B) Principal coordinate analysis (PCoA) plots of caecal bacterial community in each group. (C-D) The richness (Chao 1, C) and diversity (Shannon index, B) of caecum bacterial community in each group. (E) The relative abundance of *Lactobacillus* genus of each group. In the cases of diversity indexes and microbial taxa were analyzed by non-parametric Kruskal-Wallistest. *p < .05, **p < .01, and ***p < .001

## 4. Discussion

The results of this study demonstrate that *Limosilactobacillus fermentum* ATG-V5 has immune-enhancing effects both *in vitro* and *in vivo*. *In vitro*, ATG-V5 was found to increase the phagocytic activity of macrophages, increase the production of nitric oxide and inflammatory cytokines, and enhance the proliferation of T cells and cytolytic activity of NK cells (figure 1). These results suggest that ATG-V5 activates the cellular and humoral immune response, which contributes to eliminating intracellular pathogens.

*LimosiLactobacillus* is a newly proposed genus in the *Lactobacillus* family, and limited research is available on its immunomodulatory effects. However, a few studies have investigated the immunological properties of strains previously classified as *Lactobacillus paracasei*, which have now been reclassified as *Lactobacillus paracasei* (Kim et al., 2022; Nunez-Diaz et al., 2018). One study demonstrated that the administration of a fermented milk product containing *Limosilactobacillus* (formerly *L. paracasei*) strain Shirota to healthy adults resulted in increased NK cell activity and improved cytokine production, indicating a potential immunomodulatory effect of the strain(Rico et al., 2014). Another study investigated the immunomodulatory effects of a strain of *Limosilactobacillus fermentum in vitro* and found that it induced the production of various cytokines and enhanced the phagocytic activity of macrophages(Rodríguez-Sojo, Ruiz-Malagón, Rodríguez-Cabezas, Gálvez, & Rodríguez-Nogales, 2021).

Overall, while more research is needed to fully understand the immunomodulatory effects of *Limosilactobacillus* strains, some studies suggest that they may have potential in this area.

CPP-induced immunosuppression in mice refers to decreased in immune function due to exposure to CPP, a chemotherapy drug commonly used to treat cancer and autoimmune diseases. Specifically, CPP-induced immunosuppression can decrease T and B lymphocytes’ number and function, reduce antibody production, and impair the activation of immune cells such as macrophages and NK cells. We used a CPP-induced mouse model to evaluate the efficacy of ATG-V5-induced immunity improvement in this immunocompromised phenomenon.

*In vivo*, ATG-V5 administration was found to decrease the size and weight of the spleen, which had been increased by CPP treatment, and recover the thymus size and weight. Furthermore, ATG-V5 administration suppressed the atrophy of the white pulp in the spleen, which indicates the activation of the immune response through humoral and cell-mediated pathways. These findings suggest that ATG-V5 may have a protective effect on immune organs in mice treated with CPP.

The increase in the cytolytic activity of NK cells, the elevation of IgA, IgG, and IgM in the blood, and the increase in the number of Th1, Th2, Th17, and Treg cells suggest that ATG-V5 enhances the immune response and plays a role in the recovery of immunosuppressed mice induced by CPP.

The results of the microbiome analysis showed that the *Lactobacillus* genus, which increased due to CPP, tended to decrease under the influence of ATG-V5. A statistically significant decrease was observed when administered at high dose of ATG-V5. This result showed a similar pattern to previous studies that mentioned the relationship between *Lactobacillus* and immunoglobulin A (IgA) during immunosuppression using CPP (Mesa et al., 2020). In addition, according to Pearson’s correlation, the reference paper confirmed a significant negative correlation between immune cells and *Lactobacillus* in the CPP-treated group. These results were consistent with our experimental results, and it was confirmed that *Lactobacillus*, which was increased by CPP, decreased again in the ATG-V5 administration group. In this study, apart from the findings above, significant increases in the Ruminococcaceae and Lachnospiraceae families were also observed following ATG-V5 treatment (Supplementary Figure 1). However, these changes showed less than 1% relative abundance, and further research is necessary to explain their correlation with immune function.

In conclusion, this study suggests that *Limosilactobacillus fermentum* ATG-V5 has immunomodulatory effects both *in vitro* and *in vivo*, and may have the potential as a therapeutic agent for immune-related diseases. However, further research is needed to investigate the underlying mechanisms of the immune-enhancing effects of ATG-V5 and its potential clinical applications.

## 5. Conclusions

In conclusion, our results demonstrated that the novel strain *Limosilactobacillus fermentum* ATG-V5 exhibits immunomodulatory effects both in vitro and in vivo. Treatment with ATG-V5 activated phagocytosis, NO production, and cytokine generation in Raw264.7 cells enhancing NK cell cytolytic activity. Moreover, oral administration of ATG-V5 in a CPP-induced immunosuppressed mouse model restored the sizes of immune-related tissues and led to the recovery of NK cell activity, increased levels of immunoglobulins, and enhanced T cell distribution, as evidenced by splenocyte analysis. Furthermore, the mouse model’s intestine analysis revealed that ATG-V5 induced significant changes in the overall gut microbiota composition.

## Declaration of competing interest

All authors declare no conflicts of interest.

## Abbreviations

CPP: Cyclophosphamide
FAO: Food and Agriculture Organization
NK: Natural killer
NO: Nitric oxide
PCoA: Principal coordinate analysis
WHO: World Health Organization

---

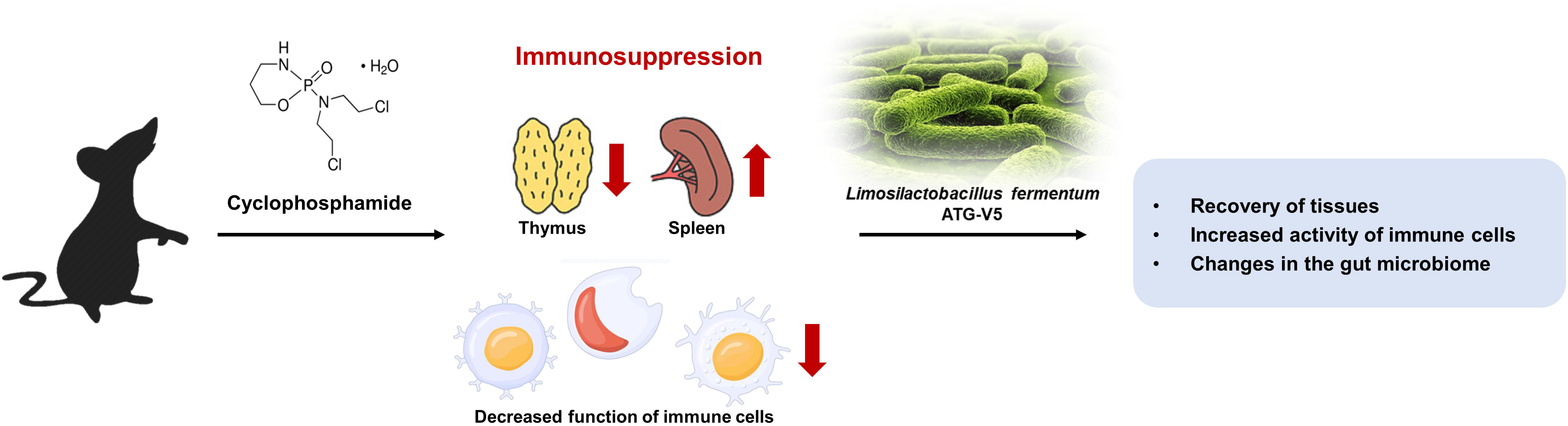

## Notes

### Competing Interest Statement

The authors have declared no competing interest.

## References

1. Baker, R. E., Mahmud, A. S., Miller, I. F., Rajeev, M., Rasambainarivo, F., Rice, B. L., . . . Metcalf, C. J. E. (2022). Infectious disease in an era of global change. Nat Rev Microbiol, 20(4), 193–205. doi: 10.1038/s41579-021-00639-z

2. Belkaid, Y., & Hand, T. W. (2014). Role of the microbiota in immunity and inflammation. Cell, 157(1), 121–141. doi: 10.1016/j.cell.2014.03.011

3. Bensinger, W. I., Martin, P. J., Storer, B., Clift, R., Forman, S. J., Negrin, R., . . . Appelbaum, F. R. (2001). Transplantation of bone marrow as compared with peripheral-blood cells from HLA-identical relatives in patients with hematologic cancers. N Engl J Med, 344(3), 175–181. doi: 10.1056/NEJM200101183440303

4. Bolyen, E., Rideout, J. R., Dillon, M. R., Bokulich, N. A., Abnet, C., Al-Ghalith, G. A., . . . Asnicar, F. (2018). QIIME 2: Reproducible, interactive, scalable, and extensible microbiome data science: PeerJ Preprints.

5. Caporaso, J. G., Lauber, C. L., Walters, W. A., Berg-Lyons, D., Huntley, J., Fierer, N., . . . Knight, R. (2012). Ultra-high-throughput microbial community analysis on the Illumina HiSeq and MiSeq platforms. [Short Communication]. The Isme Journal, 6(8), 1621. doi: 10.1038/ismej.2012.8

6. Cervantes-Barragan, L., Chai, J. N., Tianero, M. D., Di Luccia, B., Ahern, P. P., Merriman, J., . . . Colonna, M. (2017). Lactobacillus reuteri induces gut intraepithelial CD4(+)CD8alphaalpha(+) T cells. Science, 357(6353), 806–810. doi: 10.1126/science.aah5825

7. Darma, A., Athiyyah, A. F., Ranuh, R. G., Endaryanto, A., Budiono, B., & Sudarmo, S. M. (2020). Effects of probiotics on the enhancement of the innate mucosal immune response against pathogenic bacteria. Iran J Microbiol, 12(5), 445–450. doi: 10.18502/ijm.v12i5.4606

8. Emadi, A., Jones, R. J., & Brodsky, R. A. (2009). Cyclophosphamide and cancer: golden anniversary. Nat Rev Clin Oncol, 6(11), 638–647. doi: 10.1038/nrclinonc.2009.146

9. Food, F. W. J., Nations, A. O. o. t. U., & Report, W. H. O. E. C. (2001). Evaluation of health and nutritional properties of powder milk and live lactic acid bacteria (pp. 1–34): Joint FAO/WHO Expert Consultation Cordoba, Argentina.

10. Gaforio, J. J., Ortega, E., Algarra, I., Serrano, M. J., & Alvarez de Cienfuegos, G. (2002). NK cells mediate increase of phagocytic activity but not of proinflammatory cytokine (interleukin-6 [IL-6], tumor necrosis factor alpha, and IL-12) production elicited in splenic macrophages by tilorone treatment of mice during acute systemic candidiasis. Clin Diagn Lab Immunol, 9(6), 1282–1294. doi: 10.1128/cdli.9.6.1282-1294.2002

11. Geuking, M. B., & Burkhard, R. (2020). Microbial modulation of intestinal T helper cell responses and implications for disease and therapy. Mucosal Immunol, 13(6), 855–866. doi: 10.1038/s41385-020-00335-w

12. Hayase, E., & Jenq, R. R. (2021). Role of the intestinal microbiome and microbial-derived metabolites in immune checkpoint blockade immunotherapy of cancer. Genome Med, 13(1), 107. doi: 10.1186/s13073-021-00923-w

13. Hirayama, D., Iida, T., & Nakase, H. (2017). The Phagocytic Function of Macrophage-Enforcing Innate Immunity and Tissue Homeostasis. Int J Mol Sci, 19(1). doi: 10.3390/ijms19010092

14. Iddir, M., Brito, A., Dingeo, G., Fernandez Del Campo, S. S., Samouda, H., La Frano, M. R., & Bohn, T. (2020). Strengthening the Immune System and Reducing Inflammation and Oxidative Stress through Diet and Nutrition: Considerations during the COVID-19 Crisis. Nutrients, 12(6). doi: 10.3390/nu12061562

15. Jahns, L., Conrad, Z., Johnson, L. K., Whigham, L. D., Wu, D., & Claycombe-Larson, K. J. (2018). A diet high in carotenoid-rich vegetables and fruits favorably impacts inflammation status by increasing plasma concentrations of IFN-alpha2 and decreasing MIP-1beta and TNF-alpha in healthy individuals during a controlled feeding trial. Nutr Res, 52, 98–104. doi: 10.1016/j.nutres.2018.02.005

16. Jiao, Y., Wu, L., Huntington, N. D., & Zhang, X. (2020). Crosstalk Between Gut Microbiota and Innate Immunity and Its Implication in Autoimmune Diseases. Front Immunol, 11, 282. doi: 10.3389/fimmu.2020.00282

17. Kim, H., Lee, Y. S., Yu, H. Y., Kwon, M., Kim, K. K., In, G., . . . Kim, S. K. (2022). Anti-Inflammatory Effects of Limosilactobacillus fermentum KGC1601 Isolated from Panax ginseng and Its Probiotic Characteristics. Foods, 11(12). doi: 10.3390/foods11121707

18. Littman, D. R., & Rudensky, A. Y. (2010). Th17 and regulatory T cells in mediating and restraining inflammation. Cell, 140(6), 845–858. doi: 10.1016/j.cell.2010.02.021

19. Lozupone, C., Lladser, M. E., Knights, D., Stombaugh, J., & Knight, R. (2010). UniFrac: an effective distance metric for microbial community comparison. [Commentary]. The Isme Journal, 5(2), 169. doi: 10.1038/ismej.2010.133

20. Mazziotta, C., Tognon, M., Martini, F., Torreggiani, E., & Rotondo, J. C. (2023). Probiotics Mechanism of Action on Immune Cells and Beneficial Effects on Human Health. Cells, 12(1). doi: 10.3390/cells12010184

21. Mesa, D., Beirão, B. C., Balsanelli, E., Sesti, L., Caron, L. F., Cruz, L. M., & Souza, E. M. J. M. (2020). Cyclophosphamide increases lactobacillus in the intestinal microbiota in chickens. 5(4), 10.1128/msystems.00080-00020.

22. Nunez-Diaz, J. A., Fumanal, M., do Vale, A., Fernandez-Diaz, C., Morinigo, M. A., & Balebona, M. C. (2018). Transcription of IVIAT and Virulence Genes in Photobacterium damselae Subsp. piscicida Infecting Solea senegalensis. Microorganisms, 6(3). doi: 10.3390/microorganisms6030067

23. Passweg, J., & Tyndall, A. (2007). Autologous stem cell transplantation in autoimmune diseases. Semin Hematol, 44(4), 278–285. doi: 10.1053/j.seminhematol.2007.08.001

24. Rico, D. E., Marshall, E. R., Choi, J., Kaylegian, K. E., Dechow, C. D., & Harvatine, K. J. (2014). Within-milking variation in milk composition and fatty acid profile of Holstein dairy cows. J Dairy Sci, 97(7), 4259–4268. doi: 10.3168/jds.2013-7731

25. Rodríguez-Sojo, M. J., Ruiz-Malagón, A. J., Rodríguez-Cabezas, M. E., Gálvez, J., & Rodríguez-Nogales, A. (2021). Limosilactobacillus fermentum CECT5716: Mechanisms and Therapeutic Insights. Nutrients, 13(3). doi: 10.3390/nu13031016

26. Sanos, S. L., Bui, V. L., Mortha, A., Oberle, K., Heners, C., Johner, C., & Diefenbach, A. (2009). RORgammat and commensal microflora are required for the differentiation of mucosal interleukin 22-producing NKp46+ cells. Nat Immunol, 10(1), 83–91. doi: 10.1038/ni.1684

27. Shinde, R., & McGaha, T. L. (2018). The Aryl Hydrocarbon Receptor: Connecting Immunity to the Microenvironment. Trends Immunol, 39(12), 1005–1020. doi: 10.1016/j.it.2018.10.010

28. Strazar, M., Temba, G. S., Vlamakis, H., Kullaya, V. I., Lyamuya, F., Mmbaga, B. T., . . . Xavier, R. J. (2021). Gut microbiome-mediated metabolism effects on immunity in rural and urban African populations. Nat Commun, 12(1), 4845. doi: 10.1038/s41467-021-25213-2

29. van Gool, M. M. J., & van Egmond, M. (2020). IgA and FcalphaRI: Versatile Players in Homeostasis, Infection, and Autoimmunity. Immunotargets Ther, 9, 351–372. doi: 10.2147/ITT.S266242

30. van Vliet, M. J., Harmsen, H. J., de Bont, E. S., & Tissing, W. J. (2010). The role of intestinal microbiota in the development and severity of chemotherapy-induced mucositis. PLoS Pathog, 6(5), e1000879. doi: 10.1371/journal.ppat.1000879

31. van Vliet, M. J., Tissing, W. J., Dun, C. A., Meessen, N. E., Kamps, W. A., de Bont, E. S., & Harmsen, H. J. (2009). Chemotherapy treatment in pediatric patients with acute myeloid leukemia receiving antimicrobial prophylaxis leads to a relative increase of colonization with potentially pathogenic bacteria in the gut. Clin Infect Dis, 49(2), 262–270. doi: 10.1086/599346

32. Wiertsema, S. P., van Bergenhenegouwen, J., Garssen, J., & Knippels, L. M. J. (2021). The Interplay between the Gut Microbiome and the Immune System in the Context of Infectious Diseases throughout Life and the Role of Nutrition in Optimizing Treatment Strategies. Nutrients, 13(3). doi: 10.3390/nu13030886

33. Yan, F., & Polk, D. B. (2011). Probiotics and immune health. Curr Opin Gastroenterol, 27(6), 496–501. doi: 10.1097/MOG.0b013e32834baa4d

34. Yan, H., Lu, J., Wang, J., Chen, L., Wang, Y., Li, L., . . . Zhang, H. (2021). Prevention of Cyclophosphamide-Induced Immunosuppression in Mice With Traditional Chinese Medicine Xuanfei Baidu Decoction. Front Pharmacol, 12, 730567. doi: 10.3389/fphar.2021.730567

35. Yang, W., & Cong, Y. (2021). Gut microbiota-derived metabolites in the regulation of host immune responses and immune-related inflammatory diseases. Cell Mol Immunol, 18(4), 866–877. doi: 10.1038/s41423-021-00661-4

36. Yilmaz, P., Parfrey, L. W., Yarza, P., Gerken, J., Pruesse, E., Quast, C., . . . Glöckner, F. O. (2014). The SILVA and “All-species Living Tree Project (LTP)” taxonomic frameworks. Nucleic Acids Research, 42(D1), D643–D648. doi: 10.1093/nar/gkt1209

37. Ying, M., Yu, Q., Zheng, B., Wang, H., Wang, J., Chen, S., . . . Xie, M. (2020). Cultured Cordyceps sinensis polysaccharides modulate intestinal mucosal immunity and gut microbiota in cyclophosphamide-treated mice. Carbohydr Polym, 235, 115957. doi: 10.1016/j.carbpol.2020.115957

38. Zheng, D., Liwinski, T., & Elinav, E. (2020). Interaction between microbiota and immunity in health and disease. Cell Res, 30(6), 492–506. doi: 10.1038/s41422-020-0332-7

