## Supplementary figures and images for "*Limosilactobacillus fermentum* ATG-V5 stimulates the immune response by enhancing gut microbiome and metabolite in a cyclophosphamide-induced immunosuppression mouse model"

### Sup.

**Supplementary Figure 1**

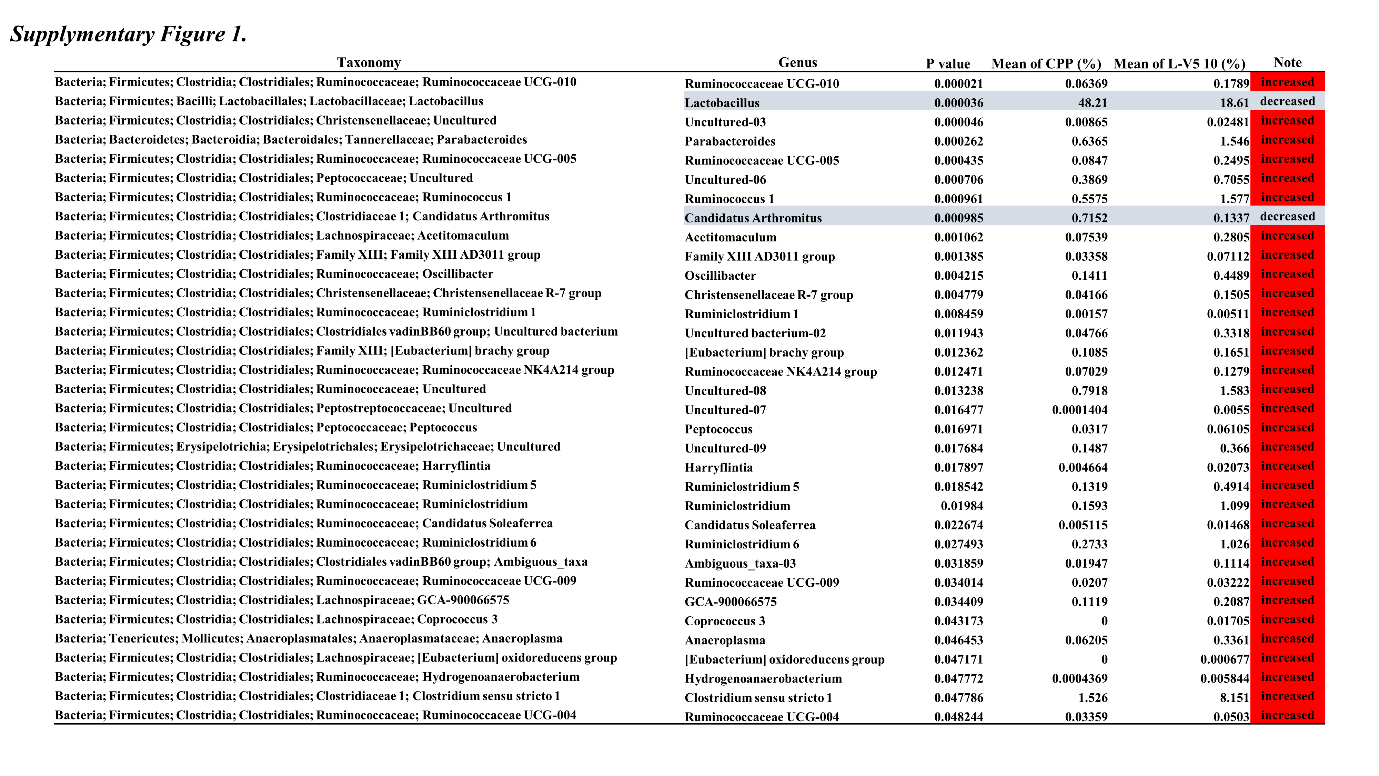
